# IgA antibodies against oxidation-specific epitopes are generated by double negative B cells and are associated with impaired lung function in fibrotic interstitial lung diseases

**DOI:** 10.64898/2025.11.28.691061

**Authors:** Eva Otoupalova, Riley T. Hannan, Pearl Raichura, Noora Batrash, Paul Dell, Monica J. Rodrigues-Jesus, Eli Zunder, Jeffrey M. Wilson, Judith A. Woodfolk, Justin J. Taylor, Shwu-Fan Ma, Andrew J. Barros, Y. Michael Shim, Imre Noth, Catherine Bonham, John S. Kim, Jeffrey M. Sturek

## Abstract

**Background:** Fibrotic interstitial lung diseases (fILD) are characterized by progressive scarring of the lung and high mortality. Oxidation-specific epitopes (OSEs) form on oxidized lipids and proteins, are abundant in lungs of patients with fILD and perpetuate lung injury and fibrosis in animal models. OSE-targeting antibodies bind to OSEs and modulate their downstream effects. The objective of this study was to determine the association of OSE-targeting antibodies with lung function and other clinical outcomes, and to characterize OSE-specific B cell phenotypes in fILD.

**Methods:** To determine the association of OSE-targeting antibodies and clinical outcomes in fILD, we measured plasma levels of OSE-specific antibodies against the malondialdehyde modified LDL mimotope P1 and anti-ApoB100 immune complex (anti-ApoB100 IC) in patients with fILD and age-matched controls. We then performed B cell phenotyping of patients with idiopathic pulmonary fibrosis (IPF) and healthy controls utilizing mass cytometry time of flight (CyTOF). To investigate effects of pro-fibrotic mediators on anti-OSE IgA antibody production, we measured B cell responses to OSEs *in vitro*.

**Results:** Cohort of 109 patients with fILD (Hypersensitivity Pneumonitis, Idiopathic Pulmonary Fibrosis, Nonspecific Interstitial Pneumonia, Connective Tissue Disease–Associated Interstitial Lung Disease) and 55 healthy donors, and a second cohort of 20 patients with IPF and 9 healthy donors were included in the study. OSE-specific antibody levels anti-P1 and anti-ApoB100 IC IgA and IgG were significantly elevated in participants with fILD compared to healthy donors (p<0.0001). Higher anti-OSE IgA was associated with worse baseline lung function, including forced vital capacity (FVC) (p<0.001) and diffusing capacity for carbon monoxide (DLCO) (p<0.01). IgA^+^CD27^-^CD21^-^CD11c^+^ double negative type 2 (DN2) B cells correlated with anti-OSE IgA levels (p<0.05) and were enriched in OSE-specific cells. DN2 B cell precursors, CXCR5^-^CD11c^+^ naive B cells and IgA^+^CD27^-^CD21^-^CD11c^-^ DN3 B cells, were significantly expanded in IPF compared to healthy donors. Both DN2 and DN3 B cells correlated with plasmablast pools in IPF patients and showed high baseline levels of IL-21 receptor, which was reduced in more differentiated cells. TGF-β combined with OSEs, but not OSEs alone, induced OSE-specific IgA production *in vitro*.

**Conclusions:** Circulating anti-OSE IgA antibodies are elevated in patients with fibrotic ILDs regardless of underlying pathology and negatively correlate with lung function. These antibodies correlated with the DN2 B cell subset which was enriched in OSE-specific cells. Phenotypic analysis of B cell subsets showed expression of markers consistent with DN to plasmablast differentiation. Finally, TGF-β promoted OSE-IgA production *in vitro*. These findings show a novel link between oxidative tissue damage, anti-OSE antibodies and DN2 B cells in fILD which may provide opportunities for new targeted therapies.

## Background

Fibrotic interstitial lung diseases (fILD) are a diverse group of autoimmune and idiopathic conditions characterized by a common terminal pathway of progressive lung scarring and respiratory failure^1,2^. The most common fILD, idiopathic pulmonary fibrosis (IPF), has an average survival of 5 years without transplant^3^. Despite intense research, therapeutic options in fILD are few and have limited efficacy. Antifibrotic medications have only modest effect on disease progression, and gastrointestinal side effects often lead to poor tolerance^4–7^. Immunosuppressive medications including corticosteroids, mycophenolate mofetil and others can improve lung function in specific types of fILD, such as systemic scleroderma-related ILD, but can be harmful in others, including IPF^8–12^. Therefore, better understanding of immunopathogenesis of fILD is urgently needed.

IPF rarely affects individuals younger then 60 years old, and average age of diagnosis is 65^3^. With advanced age, byproducts of oxidative stress including oxidatively damaged proteins, lipids and DNA, accumulate in tissues and create neo-epitopes collectively referred to as oxidation specific epitopes (OSEs)^13–15^. These conserved molecular patterns are highly immunogenic and stimulate both innate and adaptive responses. OSEs promote upregulation of inflammatory and fibrosis-associated genes, as well as secretion of pro-inflammatory and pro-fibrotic mediators including IL-6, TGF-β and MMP-9^15–17^. OSEs are abundant in bronchoalveolar lavage and lung tissue of patients with fILD^18–22^, and induce fibrotic remodeling *in vivo* in silica and bleomycin experimental models of pulmonary fibrosis^23^.

Circulating autoantibodies have been identified in several fibrotic lung diseases^24–27^, and abundant secretory IgA has been detected in the airspaces of IPF patients^28,29^. Although some of these antibodies have been linked to worse outcomes and more rapid disease progression, the antigens stimulating these responses and their significance is unclear. A subset of endogenously produced antibodies can bind to OSEs and modulate their downstream effects. These OSE-targeting antibodies are low affinity antibodies that are part of an innate-like response to exogenous and endogenous antigens^30–34^. We have previously shown that endogenous OSE-targeting IgM can attenuate lung fibrosis in an experimental bleomycin model^35^. In the current study, we tested the hypothesis that OSE-targeting antibodies are associated with survival and disease severity in patients with fILD. We observed that anti-OSE IgA was significantly elevated and correlated with higher disease severity and increased mortality in the fILD population. In addition, we also identified and characterized an OSE-specific CD27^-^CD21^-^CD11c^+^ double negative (DN) B cell population and examined its role in IPF.

## Study Design and Methods

### Ethical considerations

All data and sample collection and study procedures were approved by the Institutional Review Board for human studies at the University of Virginia (IRB numbers HSR210010, HSR200171, 15299, 20847, 20937).

### Study Subjects

For our study, we utilized banked plasma of 159 patients with fILD (IPF, hypersensitivity pneumonitis, connective tissue disease-associated ILD and non-specific interstitial pneumonia) enrolled between 2011-2015, and banked plasma and peripheral blood mononuclear cells (PBMCs) from 20 patients with IPF enrolled between 2018-2023. The diagnosis was made on the basis of multidisciplinary discussion applying contemporary ATS diagnostic criteria. Each patient was evaluated during the initial and follow-up visits and underwent PFT testing at enrollment and annually. Control group samples were from age- and sex-matched healthy donors (55 with plasma, 9 with PBMCs). Healthy donors were screened for significant history of chronic diseases and for history of any lung diseases and were excluded if either was positive. For comparison of anti-OSE antibody levels in plasma and bronchoalveolar lavage (BAL), we used paired BAL and plasma samples from patients undergoing bronchoscopy for clinical indications at UVA between 2017-2023. Details of used cohorts and workflow are specified in **Figure 1**.

**Figure 1:**
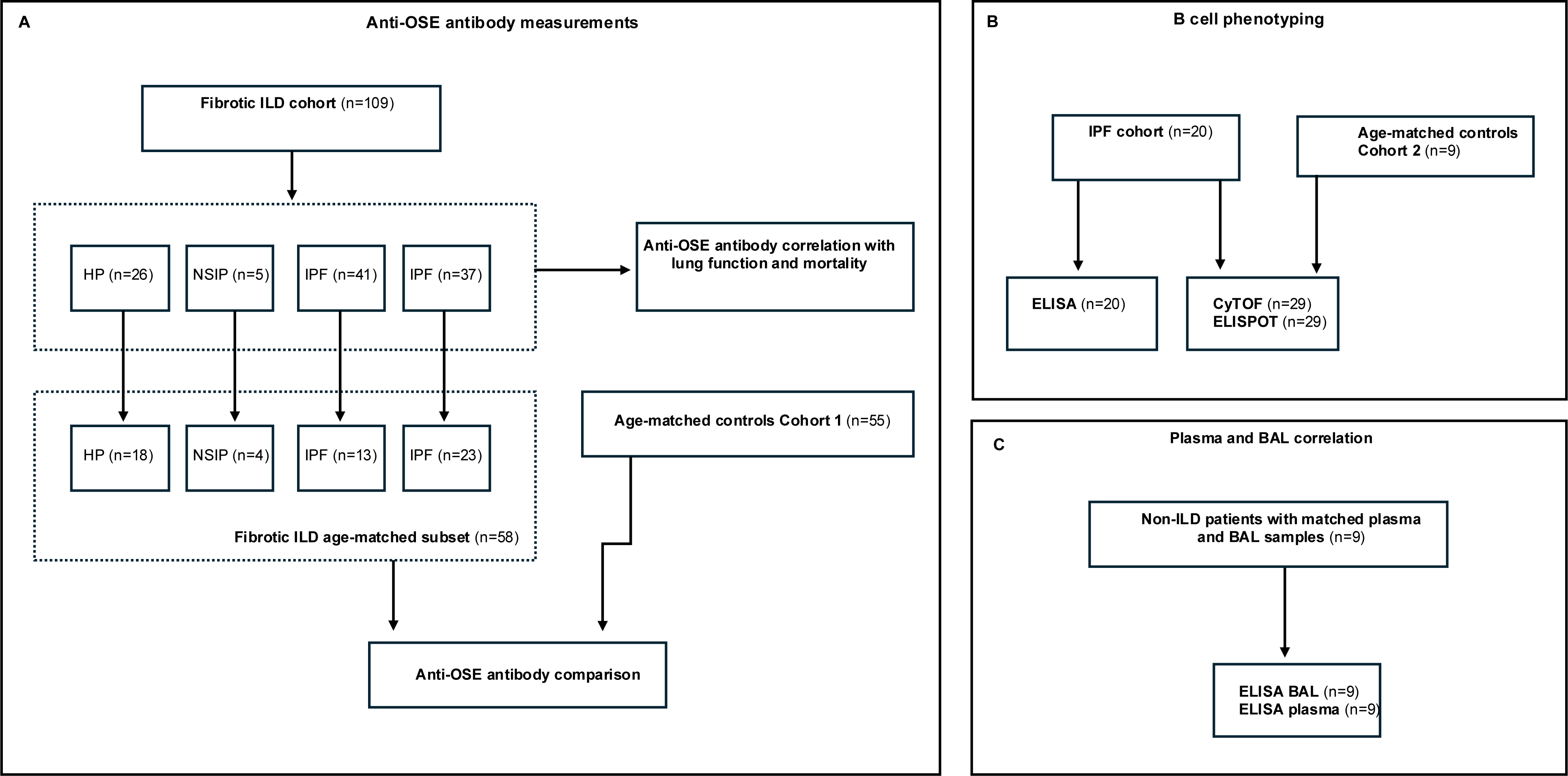
Cohort Overview. Overview of patient populations and subsets included in each experimental design: Anti-OSE antibody measurement studies with fILD cohort and fILD subset with age-matched healthy donors (A), IPF cohort for B cell phenotyping with age-matched controls (B) and non-ILD cohort for plasma/BAL correlation (C). HD=Healthy donor; fILD=fibrotic Interstitial Lung Disease; IPF=Idiopathic Pulmonary Fibrosis; HP=Hypersensitivity Pneumonitis; NSIP=Non-specific Interstitial Pneumonia; CTD-ILD=Connective Tissue Disease–associated Interstitial Lung Disease; BAL=Bronchoalveolar lavage.

### ELISA

We diluted plasma samples at a concentration of 1:10,000, 1:100,000 and 1:400,000 to measure total IgM, IgA and IgG (Jackson ImmunoResearch) by sandwich ELISA as previously described^36^. We established total antibody concentrations with human IgM, IgA and IgG (Thermo Scientific, Millipore Sigma) standard curves. To measure concentrations of anti-OSE antibodies anti-P1 and ApoB100-IC by sandwich ELISA, we coated plates with primary antigens P1 (Peptide2, P1 Mimotope, HSWTNSWMATFLGGGC) and anti-ApoB100 (Millipore Sigma Cat MAB012) and added plasma samples diluted to 1:200. We then added HRP-conjugated secondary anti-IgM, anti-IgA and anti-IgG antibodies in 1:40,000 dilution (Jackson ImmunoResearch)^37^. We measured absorbance at 450 nm. We normalized raw P1 and ApoB100-IC absorbances to internal controls to minimize batch-to-batch variability.

### ELISPOT

Human PBMCs isolated from healthy volunteers and from ILD patients were isolated and stored in liquid nitrogen as previously described^38^. For ELISPOT, CTL ImmunoSpot IgA kits (Cat# hIgA-SCE-2M/2) and IgM kits (Cat# hIgM-SCE-2M/2) were used. Cells were stimulated with TGF-β (R&D systems, 1 ng/mL) and P1 (Peptide2, P1 Mimitope, HSWTNSWMATFLGGGC, 10 mcg/mL) with BAFF (PeproTech, 400 ng/mL) containing media for 6 days and ELISPOTs were then developed per protocol (**Supplement**).

### Mass Cytometry Time of Flight

Using anti-human CD45 (HI30) conjugates available from Standard BioTools (formerly Fluidigm), we developed an eight-pick-four doublet excluding barcode to allow the multiplexed staining and acquisition of up to seventy samples simultaneously, using the methodology of Zunder et. al.^39^. CD45 based live-cell barcodes have been described previously^39–42^. Cells were barcoded using Y89, Cd106, Cd110, Cd111, Cd112, Cd113, Cd114, Cd116 as described in detail in **Supplement**. Cells were then analyzed on a Standard BioTools Cytof-2 Mass Cytometer with Helios upgrade as detailed in **Supplement**.

### Tetramer production and labeling

An OSE tetramer was generated using biotinylated cyclic OSE mimotope (P2)^37^ using the protocol from Phelps *et.al.*^43^. Biotinylated OSE mimotope P2 or scrambled control peptide (decoy) was loaded onto a streptavidin core conjugated to phycoerythrin (PE) or PE-DyLight650, respectively. 3×10^8^. 3×10^8^3-10E8 healthy donor PBMCs isolated from leukocyte reduction system (LRS) cones (StemCell 200-0093) were labeled with decoy tetramer and then P2 tetramer. B cells were then tetramer-enriched using magnetic enrichment of anti-PE-conjugated magnetic beads, followed by surface marker immunostaining. Retained portions of unenriched PBMCs were also stained. Samples were acquired on a Cytek Northern Lights analytic flow cytometer. Hierarchical gating using canonical surface markers was performed and P2-tetramer positivity defined using a P2-deficient fluorescence-minus-one (FMO) gate. Absolute and relative cell counts were calculated as necessary per donor using the original and tetramer-enriched cell counts and recorded volumes.

### Statistical Analysis

Plasma levels of total IgM, IgG, IgA and OSE-IgM, -IgG and -IgA (anti-P1, ApoB100-IC) were measured by ELISA at baseline. Primary clinical variables were forced vital capacity (FVC) and diffusing capacity of carbon monoxide (DLCO), measured at the time of sample collection, and transplant-free survival, a validated outcome for ILD. We used t-test or Mann-Whitney test to examine the difference in mean OSE-Ig levels between the fibrotic ILD and normal comparator groups using a case-control approach matched for age and gender. ANOVA or Kruskal-Wallis tests were used to examine difference in mean OSE-Ig levels by ILD diagnosis (IPF vs. Chronic HP vs. CTD-ILD). Non-parametric Spearman correlation was used to examine cross-sectional associations of baseline OSE-Ig levels, FVC and DLCO and cell populations of interest.

## Results

### Circulating anti-OSE IgA antibodies are elevated in fibrotic ILD and correlate with lung function

To test whether anti-OSE antibodies are associated with fILD, we measured plasma levels of OSE-targeting antibodies anti-P1 and anti-ApoB100 IC in patients with fibrosing ILD and age-and sex-matched healthy donors (**Table 1A and 1B**). Anti-P1 and anti-ApoB100 IC IgA and IgG antibodies were significantly elevated in 58 patients with fILD compared to 55 controls (p<0.0001) (**Figure 2A, Supplemental Figure 2A-D**). Antibody levels were similar across fILD subtypes and were not affected by immunosuppression (**Supplemental Figure 3**). Anti-OSE IgA antibody levels inversely correlated with lung function, both FVC and DLCO (**Figure 2B**). Of the IgG isotype, only anti-ApoB100 IC IgG was modestly correlated with FVC **(Supplemental Figure 2E-H**). To determine whether circulating levels of anti-OSE IgA correlate with airway levels, we measured anti-P1 antibody in paired plasma and bronchoalveolar (BAL) lavage samples of patients undergoing bronchoscopy for clinical indications. Anti-P1 IgA was detectable in BAL and strongly correlated with plasma levels (**Figure 2C**). Finally, we evaluated whether baseline anti-OSE antibody levels were associated with transplant-free survival (TFS) in patients with fILD. We stratified patients based on the range of normal anti-OSE Ig levels derived from the control cohort. Baseline anti-P1 IgA > 1.75 and anti-ApoB100 IC IgA > 0.75 relative optical density (OD) correlated with significantly lower TFS at 3 years (**Figure 2D**).

**Table 1A:** Fibrotic Interstitial Lung Disease cohort characteristics (all patients).

| Group |  | HP | NSIP | IPF | CTD-ILD | All |
| --- | --- | --- | --- | --- | --- | --- |
| Number |  | 26 | 5 | 41 | 37 | 109 |
| Age |  | 62.6 | 68 | 70.5 | 58.3 | 64.36 |
| Race | Black | 2 | 0 | 0 | 10 | 12 (10.1%) |
|  | White | 23 | 5 | 41 | 26 | 95 (88.0%) |
|  | Asian | 0 | 0 | 0 | 1 | 1 (0.91%) |
|  | Hispanic | 1 | 0 | 0 | 0 | 1 (0.91%) |
| Sex | Male | 12 | 1 | 30 | 16 | 59 (54.1%) |
|  | Female | 14 | 4 | 11 | 21 | 50 (45.9%) |

IPF=Idiopathic Pulmonary Fibrosis, HP=Hypersensitivity Pneumonitis, NSIP=Non-specific Interstitial Pneumonia, CTD-ILD=Connective Tissue Disease–Associated Interstitial Lung Disease.

**Table 1B:** Fibrotic Interstitial Lung Disease matched subset and healthy donor characteristics (age- and sex-matched cohorts).

| Group |  | HP | NSIP | IPF | CTD-ILD | All | Age-matched control |
| --- | --- | --- | --- | --- | --- | --- | --- |
| Number |  | 18 | 4 | 13 | 23 | 58 | 55 |
| Age |  | 59.7 | 66.8 | 69.3 | 51 | 59.2 | 57.3 |
| Race | Black | 1 | 0 | 0 | 8 | 9 (15.5%) | 2 (3.6%) |
|  | White | 16 | 4 | 13 | 14 | 47 (81.0%) | 48 (87.2%) |
|  | Asian | 0 | 0 | 0 | 1 | 1 (1.72%) | 4 (7.27%) |
|  | Hispanic | 1 | 0 | 0 | 0 | 1 (1.72%) | 1 (1.81%) |
| Sex | Male | 6 | 1 | 8 | 8 | 23 (39.6%) | 20 (36.3%) |
|  | Female | 12 | 3 | 5 | 15 | 35 (60.3%) | 35 (63.6%) |

IPF=Idiopathic Pulmonary Fibrosis, HP=Hypersensitivity Pneumonitis, NSIP=Non-specific Interstitial Pneumonia, CTD-ILD=Connective Tissue Disease–Associated Interstitial Lung Disease.

**Figure 2:**
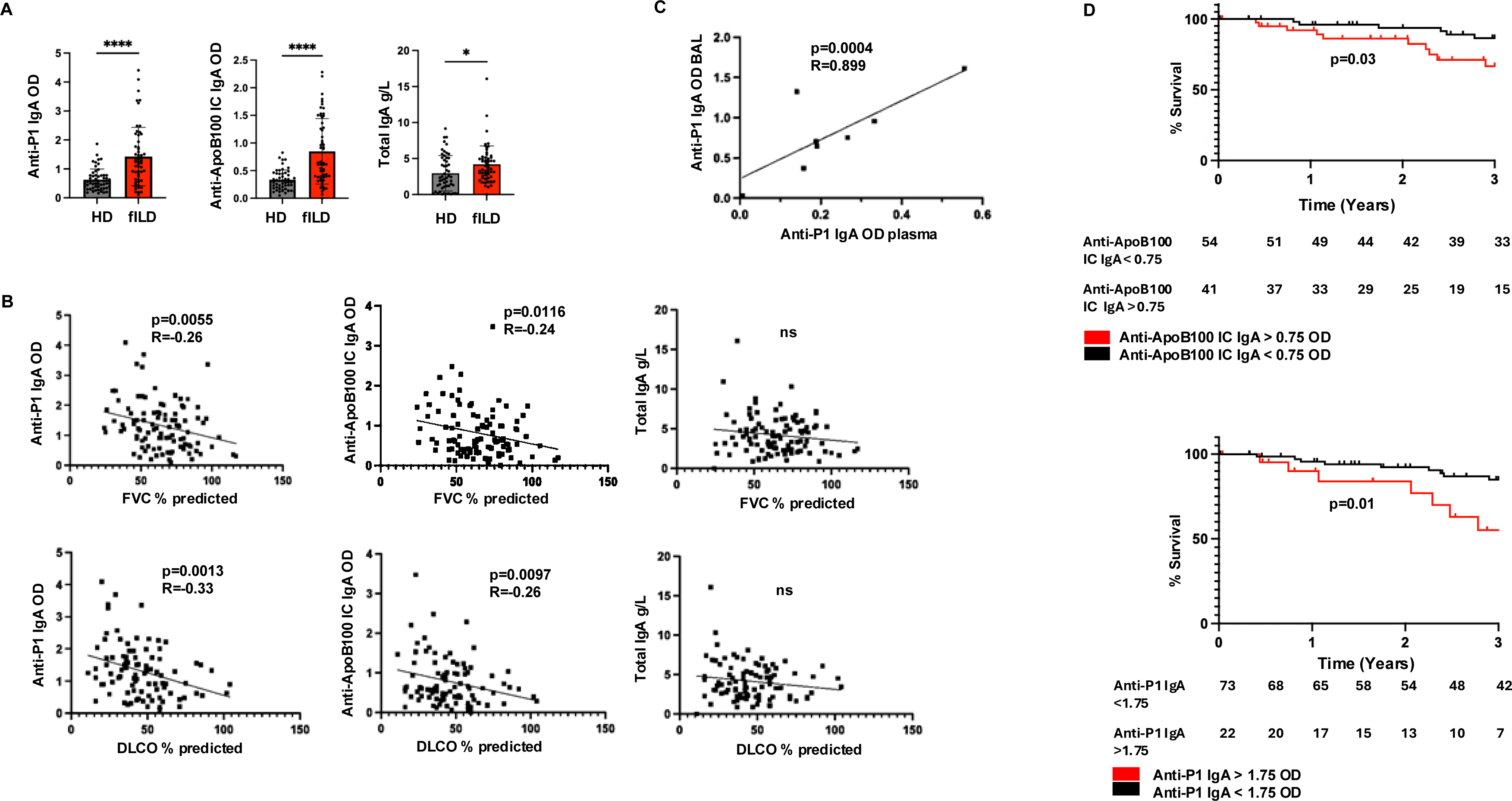
Circulating anti-OSE IgA antibodies are elevated in fibrotic ILD and correlate with lung function and mortality. Comparison of plasma anti-P1, anti-ApoB100 IC IgA and total IgA antibody levels in patients with fILD (n=58) and healthy age and sex-matched controls (n=55) (A). Correlation of plasma levels of anti-P1 IgA, andti-ApoB100 IC and total IgA (n=109) with FVC and DLCO (B). Correlation of anti-P1 IgA plasma and BAL levels in patients with non-ILD lung pathology (n=9) (C). Kaplan-Meier curves showing 3-year mortality for patients with fILD and high and low levels of anti-ApoB100 IC IgA and anti-P1 IgA (n=95) (D). All OD values were normalized to internal control. T-Test, Mann-Whitney and Spearman correlation were used for analysis. fILD=Fibrotic Interstitial Lung Disease; HD=Healthy donor; FVC=Forced vital capacity; DLCO=Diffusing lung capacity for carbon monoxide; BAL=Bronchoalveolar lavage; OD=Optical Density.

### Anti-OSE IgA antibodies positively correlate with IgA^+^CD27^-^CD11c^+^ DN2 B cells

We next analyzed surface marker expression on PBMCs from 20 patients with IPF and 9 controls with Mass Cytometry Time of Flight (CyTOF) (**Figure 3A**). We performed unsupervised hierarchical clustering of B cells using FlowSOM. We then correlated cluster frequencies with plasma OSE-IgA antibody levels (**Figure 3B; Supplemental Figure 4**). Out of 26 distinct B cell clusters, Cluster 11 strongly correlated with plasma anti-P1 and anti-ApoB IgA (p=0.017; Spearman R=0.526 and p=0.04; Spearman R=0.46, respectively). Surface marker analysis revealed high expression of CD11c, an alpha-integrin linked to atypical/age associated B cell populations, and low expression of CD27. Within Cluster 11, we additionally identified 3 distinct B cell subsets (**Figure 2C-D**): IgD^+^CXCR5^-^CD11c^+^ naive B cells, IgD^-^IgA^-^CD27^-^CD11c^+^ unswitched antigen-experienced B cells and IgD^-^IgA^+^CD27^-^CD11c^+^ switched, double negative (DN) B cells. We also observed high expression of HLA-DR, CD20, CD22 and Siglec10 and low expression of CD21 and CD24 in IgD^-^IgA^+^CD27^-^CD11c^+^ subset and Cluster 11, consistent with a DN2 population.

**Figure 3:**
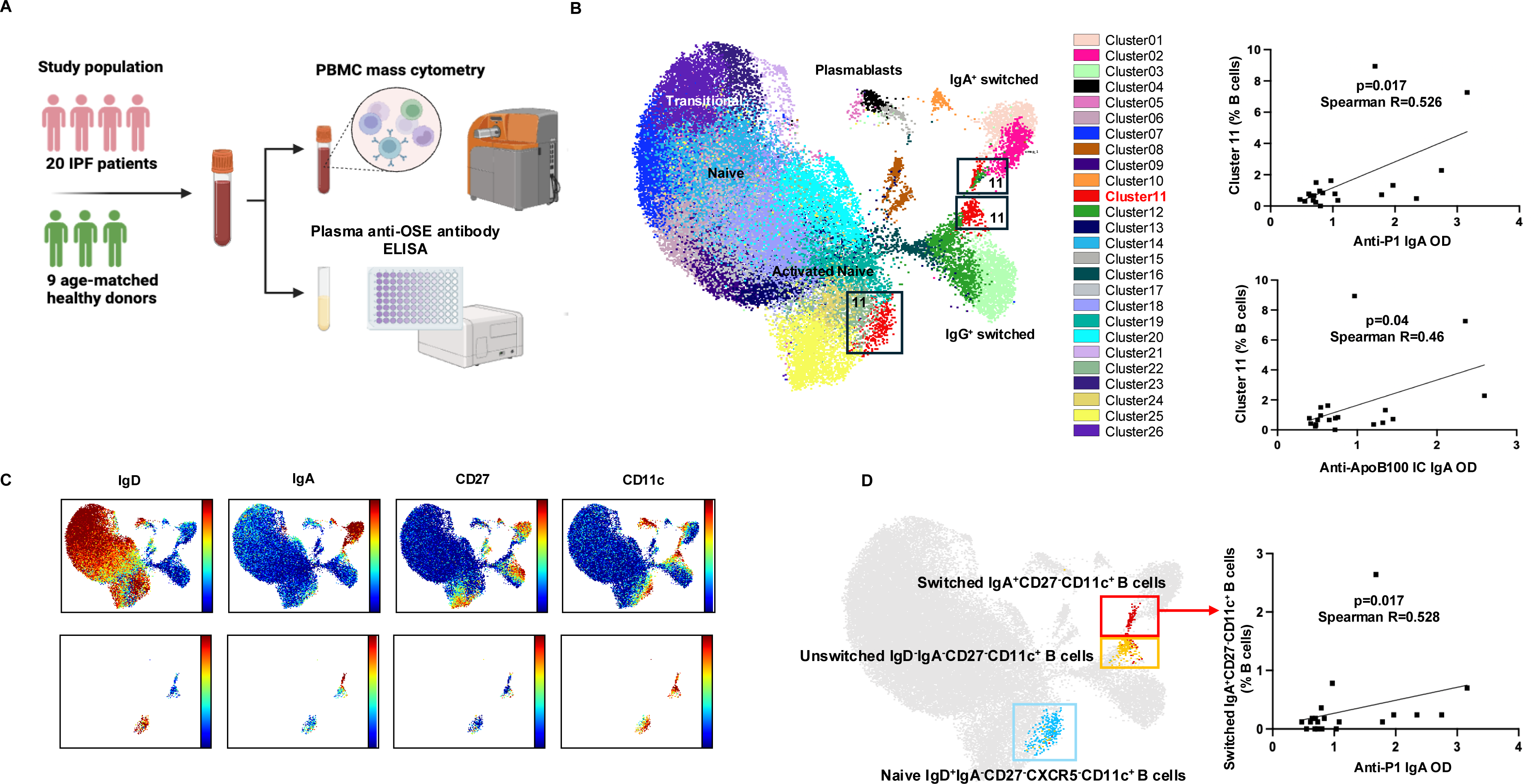
Anti-OSE IgA antibodies positively correlate with IgA^+^CD27^-^CD11c^+^ DN2 B cells. Experimental setup for Mass Cytometry Time of Flight comparing patients with IPF and age-and sex-matched healthy donors. Created in BioRender. Otoupalova, E. (2025) https://BioRender.com/gog6ivg (A). Unsupervised hierarchical clustering of B cells in patients with IPF and healthy controls showing 26 separate clusters and Spearman correlation of B cell Cluster 11 and anti-P1 IgA and anti-ApoB100 IC antibodies (p<0.05) (B). UMAP visualization of marker expression over Cluster 11 (C). UMAP visualization with overlay of naive, unswitched and switched CD27^-^CD11^+^ B cells within Cluster 11. Cluster 11 switched CD27^-^CD11^+^ B cells correlate significantly with anti-P1 OD (p<0.05) (D). All OD values were normalized to internal control. Spearman correlation was used for analysis. PBMC=Peripheral Blood Mononuclear Cells; OD=Optical Density.

### DN2 precursors are expanded in IPF, and the DN2 subset is enriched for OSE-targeting B cells

DN B cells are an antigen-experienced subset that is associated with chronic immune activation, extrafollicular antibody responses and autoimmunity. Four distinct DN B cell phenotypes have been previously described: CD21^-^CD11c^+^ B cells (DN2) are associated with extrafollicular responses and auto-antibody production, CD21^+^CD11c^-^ B cells (DN1) can differentiate into a durable switched memory B cell population, CD21^-^CD11c-B cells (DN3) are thought to be precursors of DN2s, while the role of CD21^+^CD11c^+^ (DN4) is not fully understood^44–46^. To determine alterations in DN and other B cell subsets, we used canonical gating to examine major cell populations in IPF patients and healthy donors (**Figure 4A**). Within the naive B cell population, CD11^+^CXCR5^-^ naive B cells (also described in literature as activated naive B cells^47^) and DN3 B cells were significantly expanded in IPF patients (p<0.05) (**Figure 4B**), marking a shift towards extrafollicular adaptive B cell responses. IgA^+^DN2 B cells (DN2) strongly correlated with anti-P1 IgA (p=0.003, R=0.64) and anti-ApoB100 IC IgA (p=0.002, R=0.63) (**Figure 4C**). To confirm DN2 B cell specificity for OSEs, we developed a fluorescently labeled tetramer loaded with the cyclic OSE mimotope P2 (**Figure 4D**). After B cell sample enrichment, we analyzed tetramer-positive B cell frequency in 3 healthy donors with spectral flow cytometry. While DN2 B cells formed only a small portion of total CD27^-^ and IgA^+^CD27^-^ populations, they were significantly enriched within the tetramer positive sub-populations (p<0.05), confirming that anti-OSE antibody responses are mediated through DN2 B cells.

**Figure 4:**
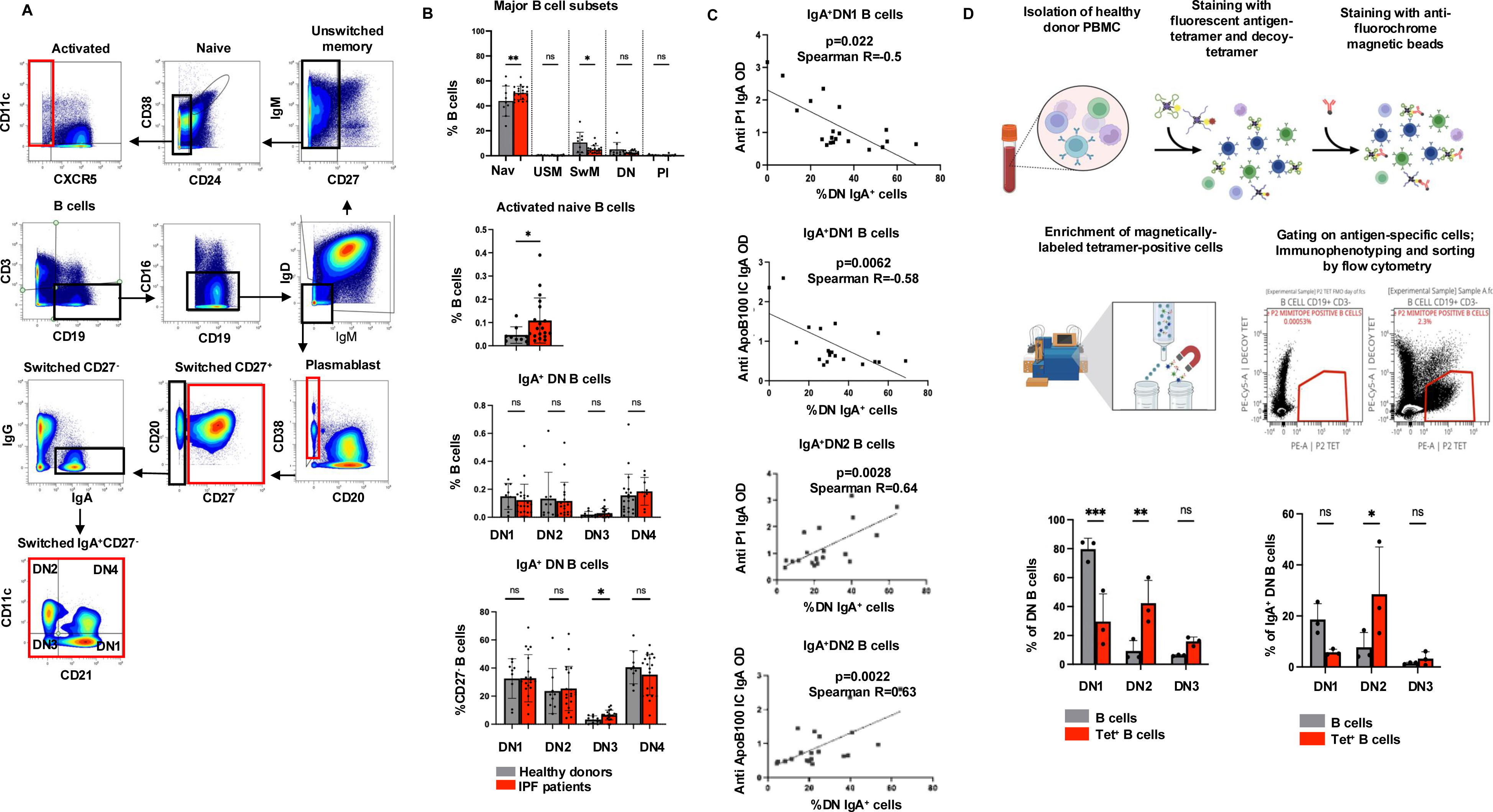
DN2 precursors are expanded in IPF, and DN2 subset is enriched for OSE-targeting B cells. Gating strategy for B cell subsets in IPF and healthy controls (A). Quantitative assessment of B cell subsets within IPF and controls (B). Association of Anti-P1 IgA and anti-ApoB100 IC IgA with frequency of double negative B cells. Spearman correlation of DN1 IgA^+^ B cells and DN2 IgA^+^ B cells and anti-P1 IgA and anti-ApoB100 IC antibodies (p<0.05) (C). Schema of tetramer positive B cell enrichment; created in BioRender. Otoupalova, E. (2025) https://BioRender.com/41zgnch, and quantification of antigen-specific B cells within double negative B cell subsets compared to unenriched B cells, summary of three independent samples from leukocyte reduction system cones (D). All OD values were normalized to internal control. Mann-Whitney, t-test, Kruskal-Wallis and ANOVA were used for analysis. PBMC= Peripheral Blood Mononuclear Cells; IPF=Idiopathic Pulmonary Fibrosis; DN=Double Negative; Nav=Naive; USM=Unswitched Memory; SwM=Switched Memory; Pl=Plasmablasts; OD=Optical Density; Tet=Tetramer.

### Reduced BAFF receptor expression on naive B cells is associated with increased extrafollicular B cell differentiation in IPF

The developmental relationships between DNs and switched memory B cell subsets remain poorly understood. We therefore used OMIQ pseudo-time PHATE analysis to assess developmental trajectories of different B cell subsets (**Figure 5A**). We identified two trajectories for antigen experienced B cells: the first one included the DN1 B cell population and a CD27^+^ switched memory population, while the second one included CXCR5^-^CD11c^+^ naive and DN2 populations.

**Figure 5:**
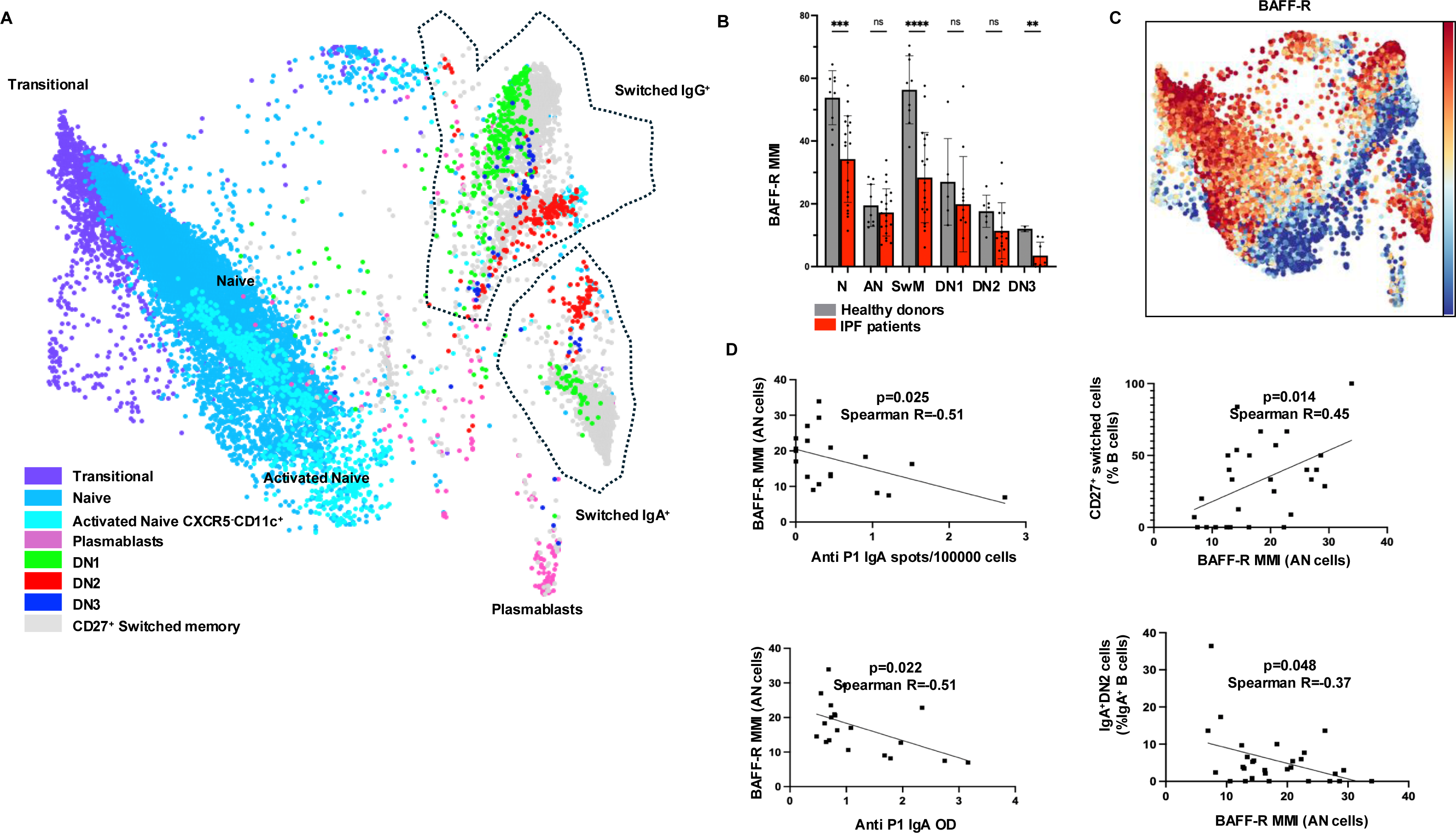
Reduced BAFF receptor expression on activated naive CXCR5^-^CD11c^+^ B cells is associated with increased extrafollicular B cell differentiation in IPF. PHATE analysis of B cell populations in IPF and control patients combined (A). Quantitative assessment of BAFF-R MMI expression on B cell subsets within IPF and controls (B). BAFF-R overlay of PHATE analysis (C). Correlation of BAFF-R MMI on CXCR5^-^CD11c^+^ naive population in IPF patients with anti-P1 IgA ELISPOT (p<0.05), anti-P1 IgA antibody (p<0.05), switched CD27^+^ (p<0.05) and IgA^+^DN2 cells (p<0.05) (D). ANOVA and Spearman correlation were used for analysis. IPF=Idiopathic pulmonary fibrosis; DN=double negative B cells; N=Naive; AN=Activated Naive; USM=Unswitched Memory; SwM=Switched memory; MMI=Mean Metal Intensity.

To determine which signaling molecules are involved in the early fate of OSE-specific B cells, we analyzed surface marker expression of naive B cell subsets. Surface BAFF receptor (CD268) levels were reduced in multiple B cell subsets in IPF patients compared to healthy controls (**Figure 5B**). BAFF expression was lower in the extrafollicular DN2 PHATE trajectory than in the CD27^+^ switched B cell trajectory (**Figure 5C**). Lower BAFF receptor expression on the CXCR5^-^CD11c^+^ activated naive subset correlated with higher IgA^+^DN2 B cell frequency (p<0.05), plasma anti-OSE IgA antibody levels (p<0.05) and anti-OSE antibody secreting cells as measured by ELISPOT (p<0.05) in IPF patients (**Figure 5D**), while higher BAFF receptor expression correlated with increased CD27^+^IgA^+^ switched memory B cell frequency. These findings support a role for decreased BAFF signaling in promoting extrafollicular B cell differentiation in IPF.

### DN2 and DN3 B cells show signs of increased differentiation into IgA^+^ plasmablasts in IPF

We next examined possible mechanisms through which DN2 cells generate more antibodies in IPF. Within the DN B cell subsets, we observed significant upregulation of IL-21 receptor (IL-21R) in IPF (**Figure 6A**). IL-21 has been shown to stimulate DN2 B cell differentiation into antibody-secreting cells. Therefore, we hypothesized that IL-21R^+^ DN2 cells in IPF are more poised to respond to IL-21 and differentiate into plasmablasts. To test this hypothesis, we evaluated expression of surface markers on IL-21R^+^ and IL-21R^-^ DN2 B cells (**Figure 6B**). IL-21R^+^ DN2 B cells demonstrated a highly activated phenotype including increased expression of CCR6, CXCR4, CXCR5, CSF1R, CD43, CD137R and CD20, consistent with an activated extrafollicular effector B cell population primed for migration and plasmablasts differentiation. Notably, we also observed an increase in early plasmablasts marker CD43 in the DN3 population, suggesting that DN3 B cells may also contribute to the plasmablast pool (**Figure 6C**). We then ran a PHATE analysis on the switched B cell subset to better delineate DN subset differentiation. While both healthy donors and IPF patients showed overlap of the DN2 B cell area with the IgG-secreting plasmablast area, IPF patients additionally showed overlap of IgA^+^DN2 and DN3 B cells with IgA-secreting plasmablasts. Notably, the plasmablast area localizing DN2 and DN3 B cells showed downregulation of IL-21 receptor (**Figure 6D**). IL-21^-^CD43^+^ DN2 B cells had the strongest correlation with anti-P1 IgA (p=0.012, Spearman R=0.54) (**Supplemental Figure 5)**. These findings suggest that IL-21 receptor signaling plays role in increased IgA^+^DN2/DN3 differentiation into plasmablasts IPF.

**Figure 6:**
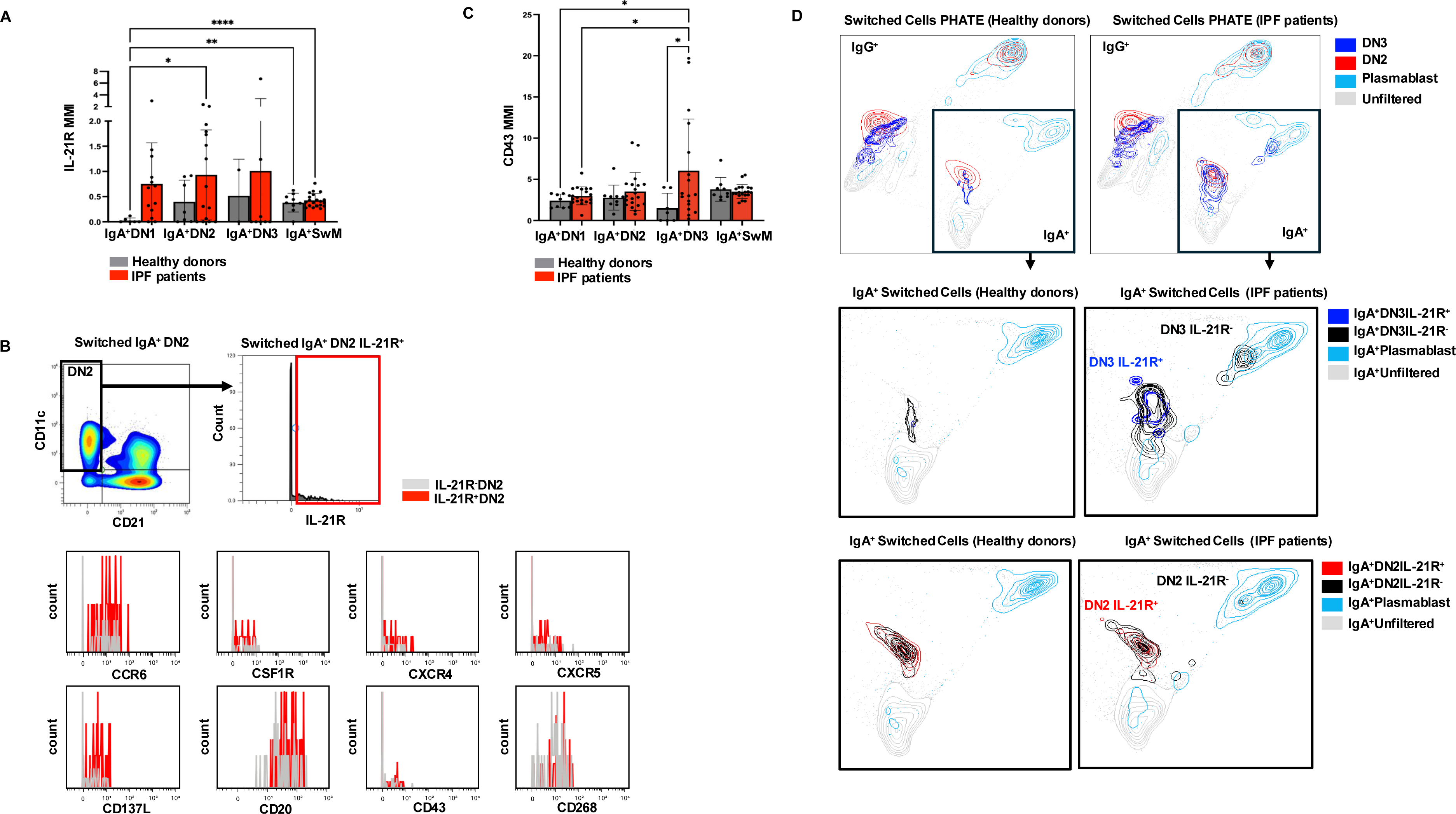
DN2 and DN3 B cells show signs of increased differentiation into IgA^+^ plasmablasts in IPF. IL-21R expression on IgA^+^ B cells (A). Expression of surface markers on switched IgA^+^DN2 IL-21R^+^ and IL-21R^-^ B cells (B). CD43 MMI expression on IgA^+^ B cells (C). PHATE analysis of switched B cells including plasmablasts with DN2 and DN3 overlays (D). ANOVA and Kruskal-Wallis was used for analysis. DN=Double Negative; SwM=Switched Memory; IL-21R=IL-21 receptor; MMI= Mean Metal Intensity.

**Figure 7:**
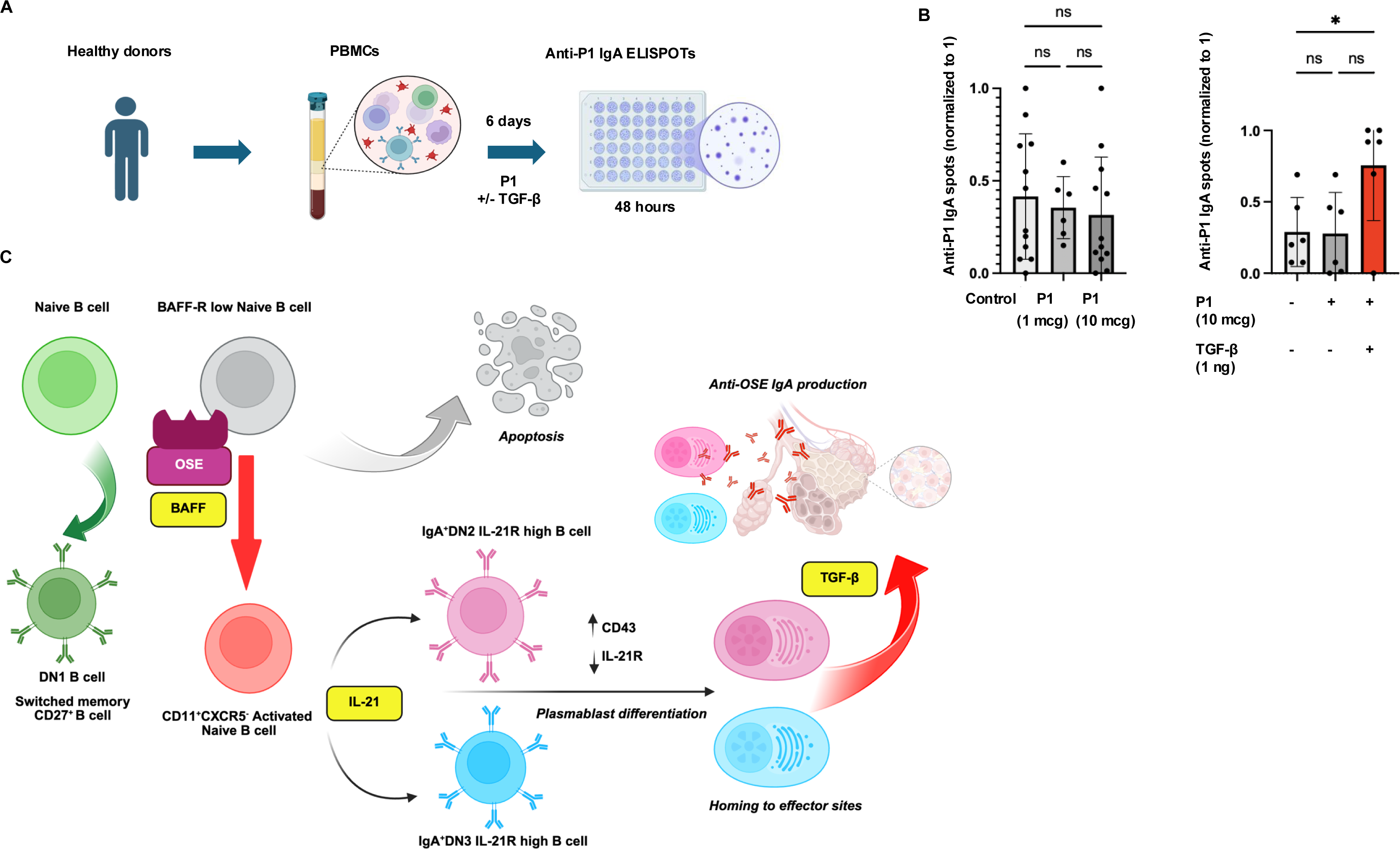
**TGF-β stimulates increased OSE-IgA production *in vitro.*** Schema of ELISPOT *in vitro* experiments. Created in BioRender. Otoupalova, E. (2025) https://BioRender.com/7dgca3x (A). ELISPOT results after *in vitro* stimulation of PBMCs, results of 4 separate experiment with 3 healthy donors (left panel) and 2 separate experiments with 3 healthy donors (right panel) normalized to 1 (B). ANOVA was used for analysis. Hypothetical model of DN cell development and immune response to OSEs. Created in BioRender. Otoupalova, E. (2025) https://BioRender.com/wqwsk8t (C). DN=Double Negative; IL-21R=IL-21 receptor; BAFF-R=BAFF receptor.

### TGF-β stimulates increased OSE-IgA production *in vitro*

TGF-β is a key profibrotic mediator in IPF. It is also known to promote class switching to IgA. To determine how TGF-β affects OSE antibody responses, we performed a series of *in vitro* experiments using PBMCs. We stimulated normal donor PBMCs with P1 antigen at 1 and 10 ng/mL for 6 days and then determined P1 specific IgA ELISPOT production (**FIgure 7A-B**). Antigen alone was not sufficient to induce secretion of anti-P1 IgA. Adding TGF-β however led to a significant increase in P1 IgA ELISPOT number.

## Discussion

In this study, we showed that circulating OSE-targeting IgA levels are significantly higher in patients with fILD compared to controls, inversely correlate with lung function, and are associated with mortality. We also identified IgA^+^DN2 B cell population as an anti-OSE antibody producing subset, and established that known DN2 precursors, CXCR5^-^CD11c^+^ naive B cells and DN3 B cells, are expanded in IPF. Additionally, we demonstrated that low BAFF receptor expression and high IL-21 receptor expression in IPF is associated with increased DN2/DN3 differentiation into plasmablasts. Finally, we demonstrated that TGF-β combined with OSE can stimulate anti-OSE IgA production *in vitro*. Together, these findings show an augmented extrafollicular IgA response against oxidation-specific epitopes in IPF that is associated with disease (**Figure 6C**).

Anti-OSE antibodies, the IgM and IgG isotypes, have been extensively studied in atherosclerosis wherein circulating anti-OSE IgM levels are associated with protection from disease while anti-OSE IgG levels are associated with increased disease^48^. To our knowledge, the current study is the first to report circulating anti-OSE IgA levels in humans in association with disease. Anti-OSE antibodies are traditionally viewed as a subset of “natural antibodies” with varying cellular origins across species. DN2 B cells have been reported to differentiate into antibody-producing cells and secrete autoantibodies^49–51^, and have recently been shown to correlate with anti-OSE IgG in atherosclerosis^52^. Here we have provided evidence connecting DN2 B cells to the anti-OSE IgA response in fILD.

Our study highlighted the association of BAFF, TGF-β and IL-21 receptor with B cell differentiation in IPF. BAFF is a factor central to B cell survival and maturation and is increased in patients with IPF^53,54^. With chronic BAFF stimulation, B cells downregulate BAFF receptors and become functionally BAFF resistant. This process leads to engagement of alternative BAFF receptors such as TACI, which shift signaling from BAFF-mediated survival towards extrafollicular differentiation^33,55–58^. However, in order for BAFF-resistant cells to avoid apoptosis, additional signals are needed, possibly in the form of chronic OSE and IL-21 stimulation. DN2 B cells with high IL-21 receptor expression have been reported in SLE^59^; such cells are prone to plasmablast differentiation and antibody secretion. Finally, TGF-β is known to promote class switching to IgA^60,61^. We observed increased expression of several tissue-homing receptors on activated IL21^+^DN2 B cells, suggesting that differentiation of plasmablasts may occur in affected lung tissue, where high local levels of TGF-β perpetuate IgA production. The possibility of plasmablasts producing antibodies in the lung has important therapeutic implications, as plasma-cell targeting therapies in lung fibrosis showed promise in animal studies^62,63^. Recent elegant spatial transcriptomic studies have further highlighted chemokine receptor signaling that may drive this process^64^. Therefore, our study emphasizes importance of recognizing different B cell functional subsets and candidate antigens that may be pathogenic in fibrotic lung diseases and facilitate development of targeted immunomodulatory therapies.

Our study has several limitations. First, although our study does show association of DN2 and anti-OSE Abs with fILD, further studies are needed to determine the mechanisms by which OSE-IgA may contribute to lung fibrosis. It is plausible that anti-OSE-IgA Abs have a pro-fibrotic effect as described previously in other studies^29,65^, however this needs to be tested in an experimental setting. Our fILD group also had significant heterogeneity, and although we found no differences across subgroups, future investigation into different ILD subtypes could provide more nuanced understanding of B cell biology. Finally, we measured antibodies and B cells only at a single time point. Serial studies might provide improved insight into B cell and antibody dynamics in fILD.

## Conclusion

Our study reports a novel association between oxidative damage and the immune response in IPF. We showed that anti-OSE IgA levels correlate with poor lung function in fILD and are increased in fILD patients. Additionally, we identified DN2 B cells as an OSE-targeting subset and demonstrated a shift towards extrafollicular immune responses and increased DN2/DN3 plasma cell differentiation in IPF.

## Acknowledgements

This work was supported in part by grants to EO (5TL1DK132771-03 from NIDDK), RTH (F32HL170760 from NHLBI), JSK (K23HL150301 and R01HL176659 from NHLBI) and JMS (Boehringer Ingelheim Discovery Award, Pulmonary Fibrosis Foundation Scholar Award, and R01HL179312 from NHLBI).

