## Supplement for "IgA antibodies against oxidation-specific epitopes are generated by double negative B cells and are associated with impaired lung function in fibrotic interstitial lung diseases"

**ELISPOT**

ELISPOT plates were pre-wetted with 70% ethanol and coated with primary antibody solution. For anti-P1 IgA and IgM detection, plates were coated with P1 antigen 10μL/mL (Peptide2, P1 Mimotope, HSWTNSWMATFLGGGC). For detection of total IgA and IgM, plates were coated with proprietary Capture Solution (CTL Immunospot) antibody mix of Igλ and Igκ 1uL/mL. Plates were then incubated in 4°C overnight. The next day, PBMCs were thawed and diluted to concentration of $10x10^6$/mL in complete media (RPMI with 10% FBS, 2mM L-glutamine, 100U/mL Penicillin, 100μg/mL Streptomycin, 8mM HEPES and 50μM 2-mercaptoethanol). Cells were then plated on the ELISPOT plates in concentration 500000/well for anti-P1 IgA and anti-P1 IgM antibody detection and 250000/well for total IgA and IgM antibody detection. Plates were incubated for 48h at 37°C in humidified incubator with 5-10% $CO_2$. Subsequently, ELISPOT plates were coated with anti-human biotin-conjugated IgM and IgA 1uL/mL and developed with CTL Tertiary SA-AP Solution and TrueBlue Substrate for 30-45 minutes per protocol. Plates were then dried, and spots counted on ImmunoSpot automatic counter and adjusted per 100000 cells.

**Mass Cytometry**

Masses used: Y89, Cd106, Cd110, Cd111, Cd112, Cd113, Cd114, Cd116. The seventy barcodes were made up and split into volumes sufficient to barcode 1-3E6 cells per aliquot and stored at -80°C until use. Cryopreserved PBMCs (90% FBS+10%DMSO) stored in vapor phase LN2 were taken immediately into a 37°C water bath, thawed until slushy, and diluted in warm RPMI+10% FBS +10 units/mL DNAse I (Sigma-Aldritch). Cells were spun down at 300xg for 10 minutes, counted and resuspended in supplemented RPMI at 5E6 cells/mL and rested for at least 30 minutes at 37°C and 5% CO2. Cells were then spun down at 300xg for 10 minutes, resuspended in 50uL of cell staining buffer (Standard BioTools or PBS+0.1% BSA) with a unique CD45 barcode, and incubated for 30 minutes at 4°C. Additional volume of staining buffer was added, cells were spun and washed 2x with staining buffer. After washing, cells were then combined into a single 15mL conical which was preabsorbed with staining buffer. Sample was spun down and resuspended in blocking buffer (cell staining buffer + 2.5% Human RealStain FC Block (Leinco). Pooled sample was then stained for 30m 4°C. Titers for the panel can be found [IN TABLE]. Sample was then spun and resuspended in a solution of 5 µM cisplatin (Standard BioTools) for 5 minutes at room temperature (25-28°C). Samples were spun and washed in cell staining media twice, and then 16% fresh, EM-grade PFA (Electron Microscopy Sciences) was added to the cell suspensions for a final concentration of 1.6% and allowed to fix for 10m at room temperature. Fixed cells were spun at 800xg for 5m, washed once, and then spun and resuspended in Maxpar Fix and Perm Buffer (Standard BioTools) with 125nM of Cell-ID Intercalator-IR (Standard BioTools) for 1hr or overnight at 4°C. Cells were spun and washed 3x in cell acquisition solution (Standard BioTools) prior to acquisition on a Standard BioTools Cytof-2 Mass Cytometer with Helios upgrade.

All FCS files had EQ beads removed and were normalized together using the Nolan lab normalizer[1]. Samples were debarcoded using the Zunder lab debarcoder[2,3]. Samples were gated using various parameters vs time, with clogs and other acquisition-interrupting events excluded by Boolean gating. Stringent event length, width, residual, offset, and center gates were used to further reduce incidence of debris and doublets[4]. Absolute thresholds for barcode separation (no less than 0.1) and mahalanobis distance (no more than 20) was used to ensure no poorly-debarcoded events were carried into subsequent analyses. If needed, further barcode quality gating was performed manually per debarcoded sample as described by Fread et al[2].

**Supplemental Figure Legends:**

**Supplemental Figure 1:** B cell Mass Cytometry Time of Flight (CyTOF) panel with markers and metals.

**Supplemental Figure 2:** Comparison of plasma anti-P1 (A) and anti-ApoB100 IC (B) IgM and plasma anti-P1 (C) and anti-ApoB100 IC (D) IgG antibody levels in patients with fILD and healthy age and sex-matched controls. Correlation of plasma levels of anti-P1 IgA with FVC (E) and DLCO (F) and anti-ApoB100 IC IgA with FVC (G) and DLCO (H). All OD values were normalized to internal control. Mann-Whitney, t-test and Spearman correlation were used for analysis. fILD=Fibrotic Interstitial Lung Disease; HD=Healthy donor; FVC=Forced vital capacity; DLCO=Diffusing lung capacity for carbon monoxide; OD=Optical Density.

**Supplemental Figure 3:** Plasma anti-P1 IgA (A), anti-ApoB100 IC IgA (B) and total IgA levels (C) in patients with different ILD sub-types. Plasma anti-P1 IgA (D) and anti-ApoB100 IC IgA (E) and total IgA levels (F) in patients with and without immunosuppression. All OD values were normalized to internal control. Mann-Whitney, t-test, ANOVA and Kruskal-Wallis were used for analysis. IPF=Idiopathic Pulmonary Fibrosis; HP=Hypersensitivity Pneumonitis; NSIP=Non-specific Interstitial Pneumonia; CTD-ILD=Connective Tissue Disease–Associated Interstitial Lung Disease; IS=Immunosuppression.

**Supplemental Figure 4:** Clustered heatmap from unsupervised hierarchical clustering, with Cluster 11 highlighted. Darker colors indicate higher median metal intensity.

**Supplemental Figure 5:** Correlation of $IgA^{+}DN2$ $IL21^{-}CD43^{+}$ cell frequency with anti-P1 IgA OD. Spearman correlation used for analysis. OD values normalized to internal standard. DN=Double negative; OD=Optical Density.

**Supplemental Figure 1**

| Marker | Metal | Marker | Metal |
|---|---|---|---|
| **CCR6** | 141 Pr | **IgG** | 160 Gd |
| **CD19** | 142 Nd | **TLR4** | 161 Di |
| **CSF1R** | 143 Nd | **CD69** | 162 Dy |
| **CD38** | 144 Nd | **CXCR3** | 163Dy |
| **CD81** | 145 Nd | **CXCR5** | 164Dy |
| **IgD** | 146 Nd | **CD40** | 165 Ho |
| **CD11c** | 147 Sm | **Siglec 10** | 166 Er |
| **IgA** | 148 Nd | **CD27** | 167 Er |
| **CD200** | 149 Sm | **CCR7** | 168 Er |
| **CD43** | 150 Nd | **CD24** | 169 Tm |
| **CD14** | 151 Eu | **CD3** | 170 Er |
| **CD95** | 152 Sm | **CD20** | 171 Yb |
| **CCR2/CD 192** | 153 Eu | **IgM** | 172 Yb |
| **IL21R** | 154Sm | **CD137** | 173 Yb |
| **CD268 (BAFFR)** | 155 Gd | **HLA-DR** | 174 Yb |
| **CD86** | 156 Gd | **CXCR4** | 175 Lu |
| **CD137L** | 158 Gd | **CD21** | 176 Yb |
| **CD22** | 159 Tb | **CD16** | 209Bi |

### Supplemental Figure 2

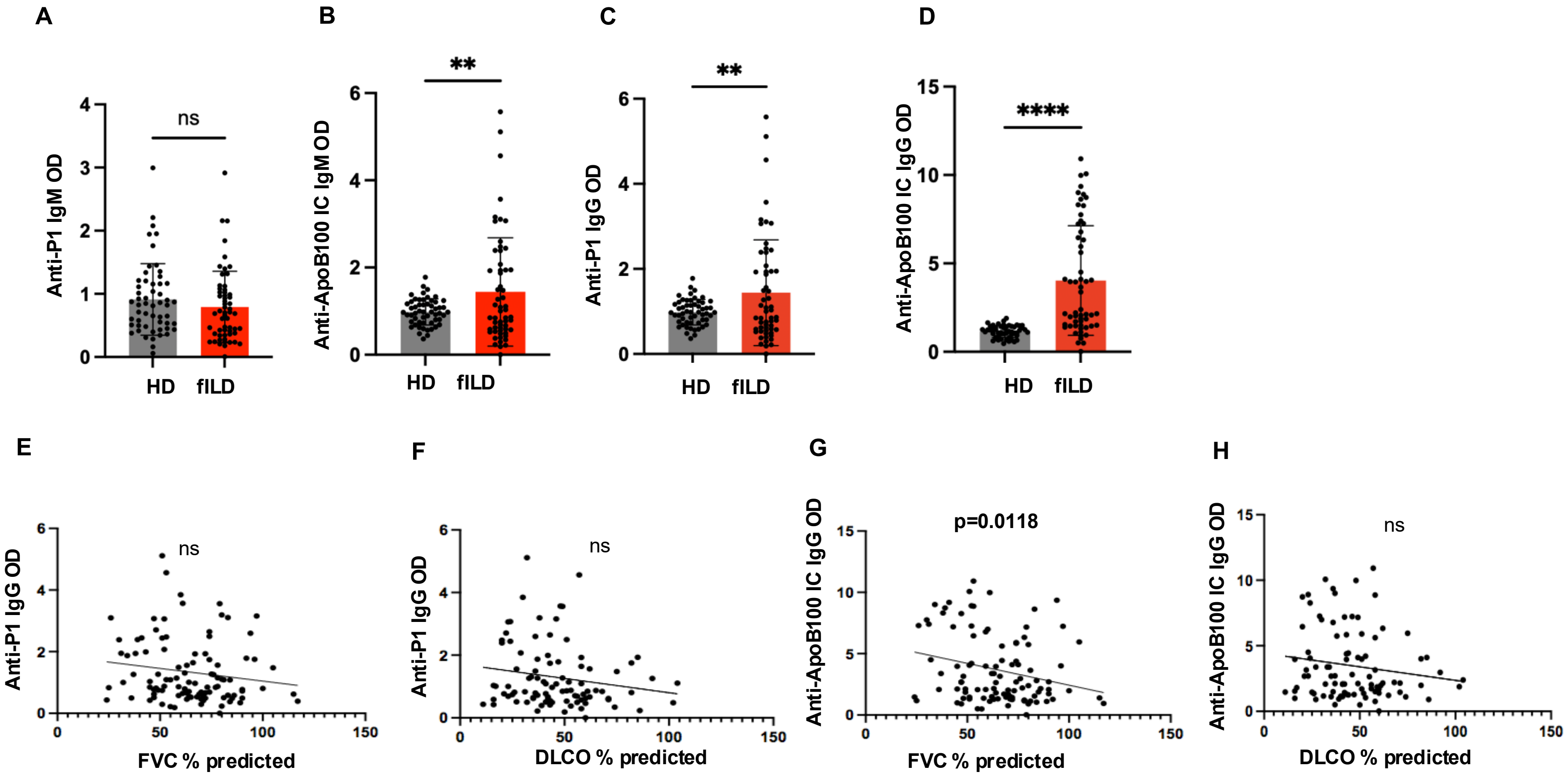

**Supplemental Figure 3**

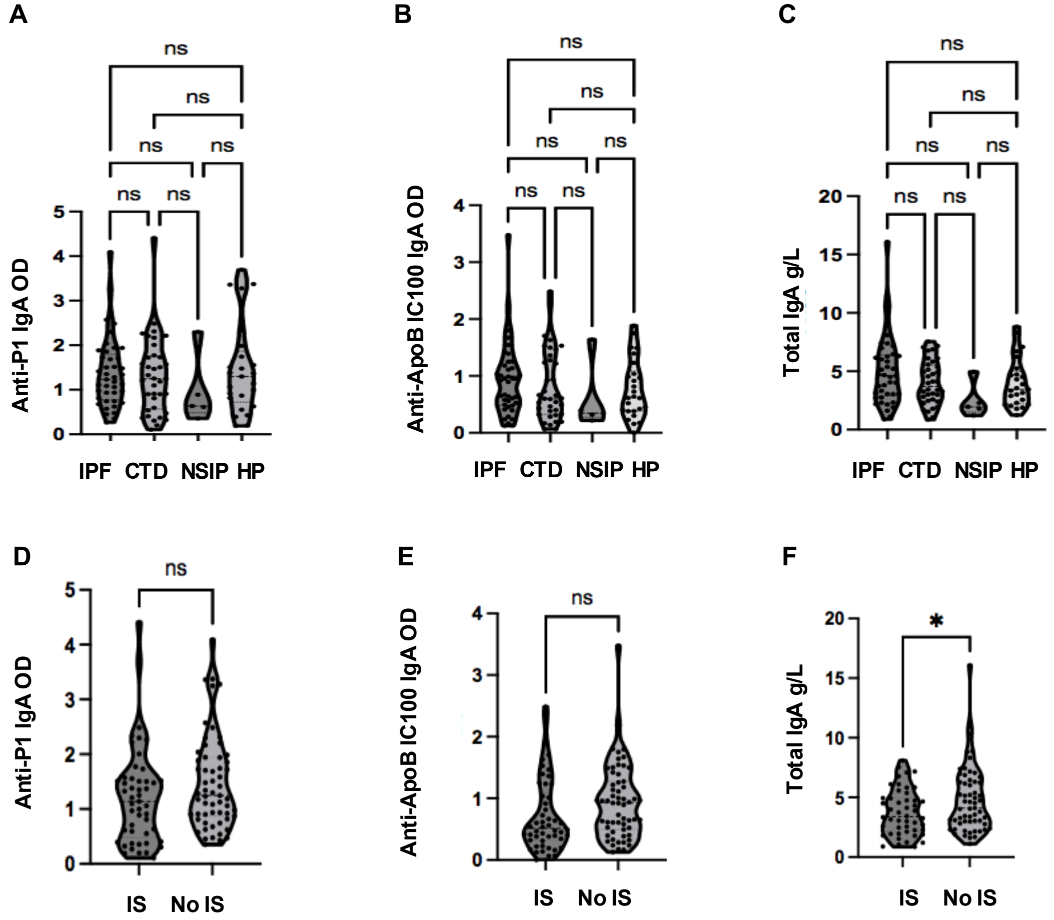

#### Supplemental Figure 4

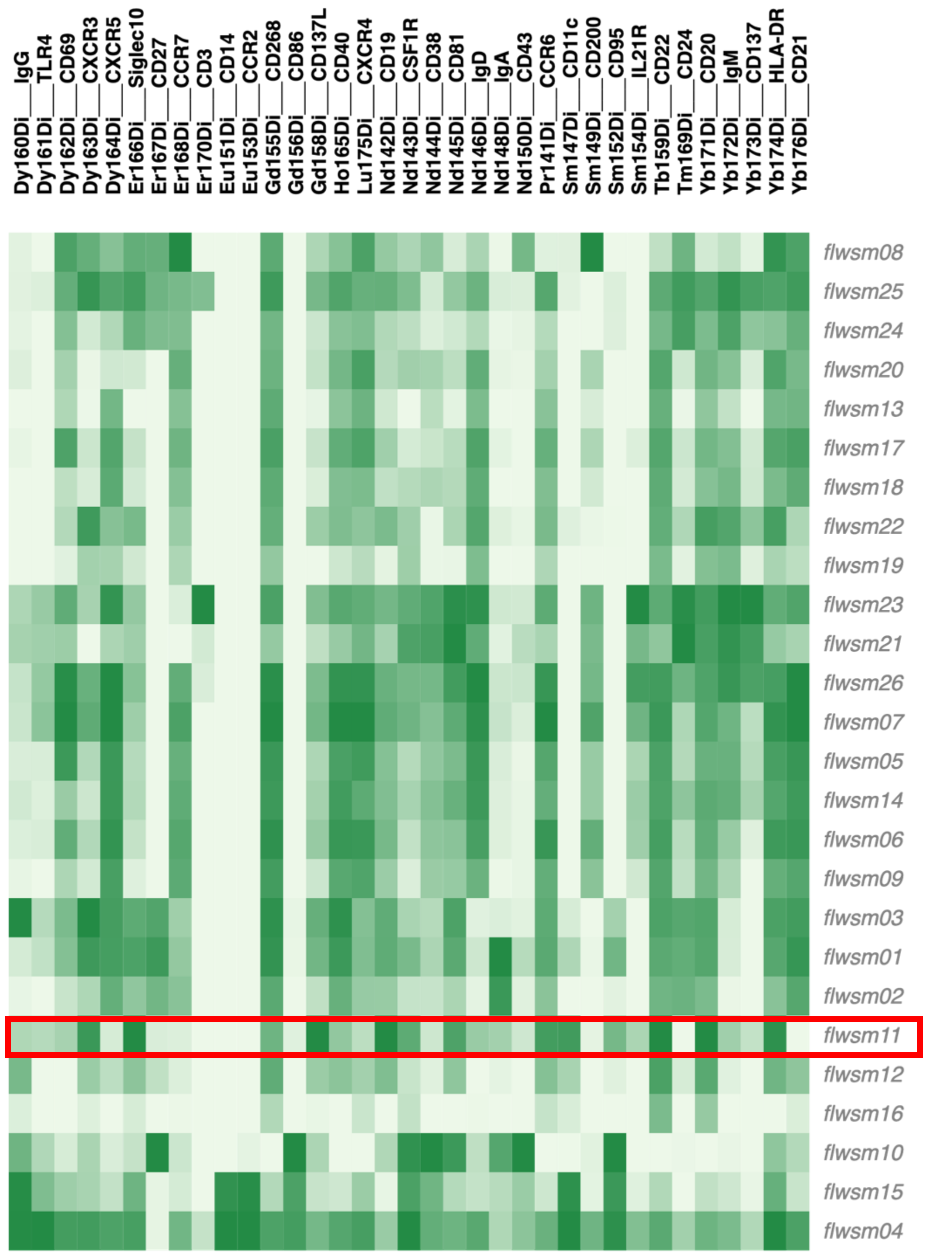

**Supplemental Figure 5**

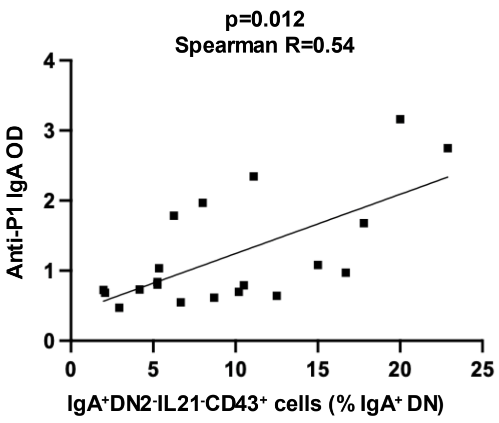
